# Subordinate effect of −21M HLA-B dimorphism on NK cell repertoire diversity and function in HIV-1 infected individuals of African origin

**DOI:** 10.1101/786392

**Authors:** Elia Moreno Cubero, Ane Ogbe, Myron S. Cohen, Barton F. Haynes, Persephone Borrow, Dimitra Peppa

## Abstract

Natural Killer (NK) cells play an important role in antiviral defence and their potent effector function identifies them as key candidates for immunotherapeutic interventions in chronic viral infections. Their remarkable functional agility is achieved by virtue of a wide array of germline encoded inhibitory and activating receptors ensuring a self-tolerant and tunable repertoire. NK cell diversity is generated by a combination of factors including genetic determinants and infections/environmental factors, which together shape the NK cell pool and functional potential. Recently a genetic polymorphism at position −21 of HLA-B, which influences the supply of HLA-E binding peptides and availability of HLA-E for recognition by the inhibitory NK cell receptor NKG2A, was shown to have a marked influence on NK cell functionality in healthy human cytomegalovirus (HCMV) seronegative Caucasian individuals. In this study, −21 methionine (M)-expressing alleles supplying HLA-E binding peptides were largely poor ligands for inhibitory killer immunoglobulin-like receptors (KIRs), and a bias to NKG2A-mediated education of functionally-potent NK cells was observed. Here, we investigated the effect of this polymorphism on the phenotype and functional capacity of NK cells in a cohort of African individuals with human immunodeficiency virus type 1 (HIV-1)/HCMV co-infection. A similarly profound influence of dimorphism at position −21 of HLA-B on NK cells was not evident in these subjects. They predominantly expressed African specific HLA-B and −C alleles that contribute a distinct supply of NKG2A and KIR ligands, and these genetic differences were compounded by the marked effect of HIV/HCMV coinfection on NK cell differentiation. Together, these factors resulted in a lack of correlation of the HLA-B −21 polymorphism with surface abundance of HLA-E and loss of the NK cell functional advantage in subjects with −21M HLA-B alleles. Instead our data suggest that during HIV/HCMV co-infection exposure of NK cells to an environment that displays altered HLA-E ligands drives adaptive NKG2C+ NK cell expansions influencing effector responses. Increased efforts to understand how NK cells are functionally calibrated to self-HLA during chronic viral infections will pave the way to developing targeted therapeutic interventions to overcome the current barriers to enhancing immune-based antiviral control.

## Introduction

There is a pressing need to better characterize and harness the immune response in order to develop efficacious immune-based strategies to supplement current therapeutic approaches for a ‘functional’ cure in chronic viral infections. Natural Killer (NK) cells have the potential to respond to viruses direct effectors and can edit adaptive immunity influencing the outcome of viral infections (1). More recently their capacity to develop adaptive or memory-like features in the setting of infection has been highlighted (2). A number of studies, both epidemiological and functional, have provided evidence for the important role of NK cells in human immunodeficiency virus type 1 (HIV-1) viral control and protection from acquiring new infection (3).

In order for NK cells to gain functional competence they are required to be ‘licensed’ or educated, a process that refines their levels of responsiveness(4). Traditionally this was ascribed to the presence of inhibitory killer immunoglobulin-like receptor (KIR) - human leukocyte antigen (HLA) class I pairs. However, recent evidence suggests that NK cells can be educated through the older and more conserved inhibitory receptor CD94/NKG2A, which recognizes HLA-E complexed with a peptide derived from the leader sequence of HLA-A, B or C alleles as well as HLA-G (5). HLA-E has little polymorphism and its levels of expression are influenced by peptide ligand availability. Where as HLA-A and HLA-C allotypes are fixed for Methionine (−21M), HLA-B contains a polymorphism that can encode either Methionine (−21M), which gives rise to functional HLA-E binding peptides, or Threonine (−21T) at this position, which does not bind effectively to HLA-E. The resultant HLA-B −21M/T variation defines different sets of haplotypes with −21M biasing toward NKG2A NK cell education, which has been shown to be associated with superior NK function in healthy HCMV seronegative adults, and −21T promoting KIR mediated education (6). The reported linkage disequilibrium (LD) in Eurasian populations between HLA-B −21M and HLA-B Bw6/HLA-C1, which interact poorly with KIRs, further decreases their potential to mediate NK cell education through KIR engagement. In contrast, −21T HLA-B haplotypes in various combinations with Bw4, C1 and C2 enhance education via KIRs. Interestingly haplotypes combining HLA-C2 and −21M HLA-B are more frequently found in Africa in combination with HLA-C allotypest hat promote HLA-E expression poorly (7). The dimorphism at position −21 of HLA-B (M/M genotype) has been associated with increased susceptibility to HIV-1 infection (8). Notably, the dimorphism influences NK cell cytolysis of HIV-infected CD4 T cells and macrophages *in vitro,* with −21T enhancing cytolysis compared to −21M, suggesting that the more educated NKG2A+ NK cell s of M/M donors may be less effective in responding to HIV-1 (9). In light of this, the beneficial effects of Bw4+ HLA-B homozygosity in controlling HIV-1 viraemia could be re-interpreted in terms of a mechanism involving recognition of HLA-E by NKG2A+ NK cells of T/T donors(10). Recently HLA-B haplotypes that favour education via NKG2A were also found to exacerbate the detrimental effect of high HLA-A on HIV-1 control through impaired killing of HIV infected target cells (11). However, this effect of HLA-A expression on HIV-1 viraemia was less pronounced in individuals of African descent, possibly reflecting the distinct frequencies of HLA haplotypes present in these populations (11).

To date, the phenotypic and functional effects of HLA-B −21 dimorphism on NK cells have not been assessed in the context of HCMV seropositive individuals or in HIV infected cohorts, where HCMV co-infection is almost universal (12). We have recently demonstrated the potent effect of HCMV co-infection in shaping the NK cell repertoire during chronic HIV-1 infection, leading to an accelerated differentiation and adaptive reconfiguration of the NK cell compartment and expansion of an NK cell subset expressing NKG2C, the activating counterpart of NKG2A that also binds to HLA-E (recognizing HLA-E bound to HLA class Ia signal sequence peptides with lower affinity than NKG2A) (13) (14) (15). The relevance of HLA-E/NKG2C interactions has been well demonstrated in driving adaptive NK cell expansions and more recently a highly specific recognition of certain HCMV-encoded HLA-E presented peptides was elegantly shown (16) (17). A rare UL40 peptide, identical to the HLA-E-binding peptide in the HLA-G signal sequence, was found to trigger optimal NK stimulation and to have functional consequences, influencing NK cell antibody dependent cellular cytotoxicity (ADCC) responses(17).

It remains unclear how the presence of this polymorphism and changes in the HLA-E ligandome during infection and inflammation affect NK cell phenotypic and functional diversity in heterogenous populations with HIV-1 infection and high levels of HCMV co-infection. To further explore this, in the current study we investigated whether the HLA-B −21 dimorphism leads to a NK cell functional dichotomy in an African cohort co-infected with HIV/HCMV.

## Materials and methods

### Study Subjects

Cross-sectional analysis was performed on peripheral blood mononuclear cells (PBMCs) cryopreserved from chronically HIV-1 infected HCMV seropositive females recruited into the Centre for HIV/AIDS Vaccine Immunology (CHAVl)00l study at clinical sites in Africa. The CHAVI001 study was approved by the Duke Medicine and National Institutes of Health Institutional Review Boards as well as the ethics boards of the local sites. All study participants gave written informed consent and were hepatitis C virus antibody negative and hepatitis B surface antigen (HBsAg) antibody negative. Human cytomegalovirus (HCMV) infection status was determined by HCMV IgG enzyme-linked immunosorbent assay (ELISA) (BioKit) on stored plasma samples. HLA class I genotyping of the study donors to 2-digit allele resolution was performed by ProImmune (Oxford, UK) by PCR analysis of DNA extracted from donor PBMC. HLA-A expression model estimates (z-score) were inferred as previously described (11). The subject characteristics, HLA class I genotypes and distribution of HLA-B −21M and −21T among HLA-B groups are summarised in **Suppl. Table 1.**

### Monoclonal Antibodies and Flow Cytometry analysis

For flow cytometric analysis, cryopreserved PBMC were thawed, washed in phosphate-buffered saline (PBS), and surface stained at 4°C for 20 min with saturating concentrations of different combinations of the following antibodies in the presence of fixable live/dead stain (Invitrogen): CD14 BV510, CD19 BV510, CD56 PE Dazzle, CD3 BV650, CD16 PERCP or CD16 BV711, HLA-E PE (3D12) (Biolegend), CD4-eFluor 780, CD8 Alexa700 (eBioscience), HLA-C PE (DT-9) (BD Biosciences), NKG2A Pe-Cy7, KIR2DL2 APC CD158b1/b2.j APC (Beckman Coulter), NKG2C PE or NKG2C Alexa 700, KIR2DL1/2DS5 APC IgG1 [CD158a], KIR3DL2 APC (R&D systems), CD57 BV421 or CD57 FITC (BD Biosciences), KIR3DL1 APC [CD158e1] (Miltenyi). For the detection of intracellular antigens cells were fixed, permeabilized and stained for IFN-γ BV421(BD Biosciences) and FcєRl-γ-FITC (Millipore). The antibody against PLZF CF594 (BD Biosciences) was used for intranuclear antigen detection utilising the Foxp3 intranuclear staining buffer kit (eBioscience) according to the manufacturer’s instructions. Samples were acquired on a BD Fortessa X20 using BD FACSDiva8.0 (BD Bioscience). Data were analyzed using FlowJo 10 (TreeStar) and stochastic neighbour embedding (SNE) analysis was performed using the mrc.cytobank platform. The FCS file concatenation tool was used for concatenating multiple FCS files into a single FCS file prior to uploading the files to cytobank.

### Functional ADCC assay

For analysis of NK cell mediated ADCC responses, RAJI cells (1-2×106 cells/mL) were coated with anti-CD20 or murine immunoglobulin G (lgG) (lnvivoGen) at 2.5 ug/ml for 30 min. Subsequently RAJI cells were washed and then mixed with PBMCs in a V bottom 96-well plate at a 10:1 ratio and incubated for 6 hours 37°C in the presence of CD107a-APC-Cy7 antibody (BD Biosciences, Cowley, U.K.). GolgiStop (containing Monensin, 1/1500 concentration, BD Biosciences) and GolgiPlug (containing brefeldin A, 1/1000 final concentration, BD Biosciences) were added for the last 5 hours of culture. Following incubation cells were washed and stained for extracellular receptors prior to permeabilization and intracellular staining for IFN-γ. Boolean gating analysis was used to analyse CD107a and IFN-γ production in CD56^dim^ NK cell subpopulations expressing CD57, NKG2A and KIRs and combinations thereof.

### Soluble HLA-G measurement

Soluble HLA-G1/G5 was measured in plasma by ELISA using a BioVendor-EXBIO kit according to the manufacturer’s instructions.

### Data analysis

Prism 7 (GraphPad Software) was used for all statistical analysis as follows: the Mann-Whitney U-test or Student’s t test were used for single comparisons of independent groups, the Wilcoxon-test was used to compare two paired groups and the Kruskal-Wallis with Dunn’s multiple comparison test was used to compare three unpaired sample groups. The nonparametric Spearman test was used for correlation analysis. SPICE analysis was performed in SPICE version 6. *p-value⍰<⍰0.05, **p- value⍰<⍰0.01, ***p-value⍰<⍰0.001, ****p-value⍰<⍰0.0001.

## Results

### Haplotypes combining HLA-C2 and −21M HLA-B are common in African populations and the HLA-B −21M dimorphism does not significantly impact on surface HLA-E expression

To explore the effects of the HLA-B dimorphism in a non-Caucasian population, we initially analyzed HLA haplotypes and examined the segregation of HLA-C allotypes and −21 HLA-B alleles in a cohort of viraemic age-matched HIV-1 infected HCMV-seropositive African females, representing the three key −21 HLA-B genotypes: −21M/ M homozygotes, −21M/T heterozygotes and −21T/T homozygotes **(Fig. 1A).** There were no significant differences in the HIV-1 viral load levels between the three groups **(Suppl. Table 1).** In contrast to Eurasian populations, which have an effective exclusion of −21M HLA-B from haplotypes encoding HLA-C2, this segregaiton was not evident in this cohort (**Fig. 1A)**, in keeping with the presence of African specific alleles, B*42:01-C*17:01 and B*81:01-C*18:01 in the M/M group **(Suppl. Table 1.).** Such haplotypes combining HLA-C2 with −21M HLA-B provide both a C2 allele, a stronger KIR ligand than C1, and an HLA-E ligand for NKG2A. HLA-B −21M alleles did not encode HLA-B Bw4 in our cohort, in line with data derived from larger population analysis (6) but interestingly, a high proportion of the subjects with −21T HLA-B alleles(9 out of 13 subjects) also did not encode HLA-B Bw4, which functions as a KIR ligand **(Suppl. Table 1).** The subsets of HLA haplotypes in the study groups defined by the presence of −21M HLA-B in various combination with HLA-Cl and C2 could therefore result in the availability of KIR ligands differentially supplying HLA-E-binding peptides to form NKG2A ligands being distinct from that in Caucasian populations, with consequences for NK cell education.

**Fig. 1.**
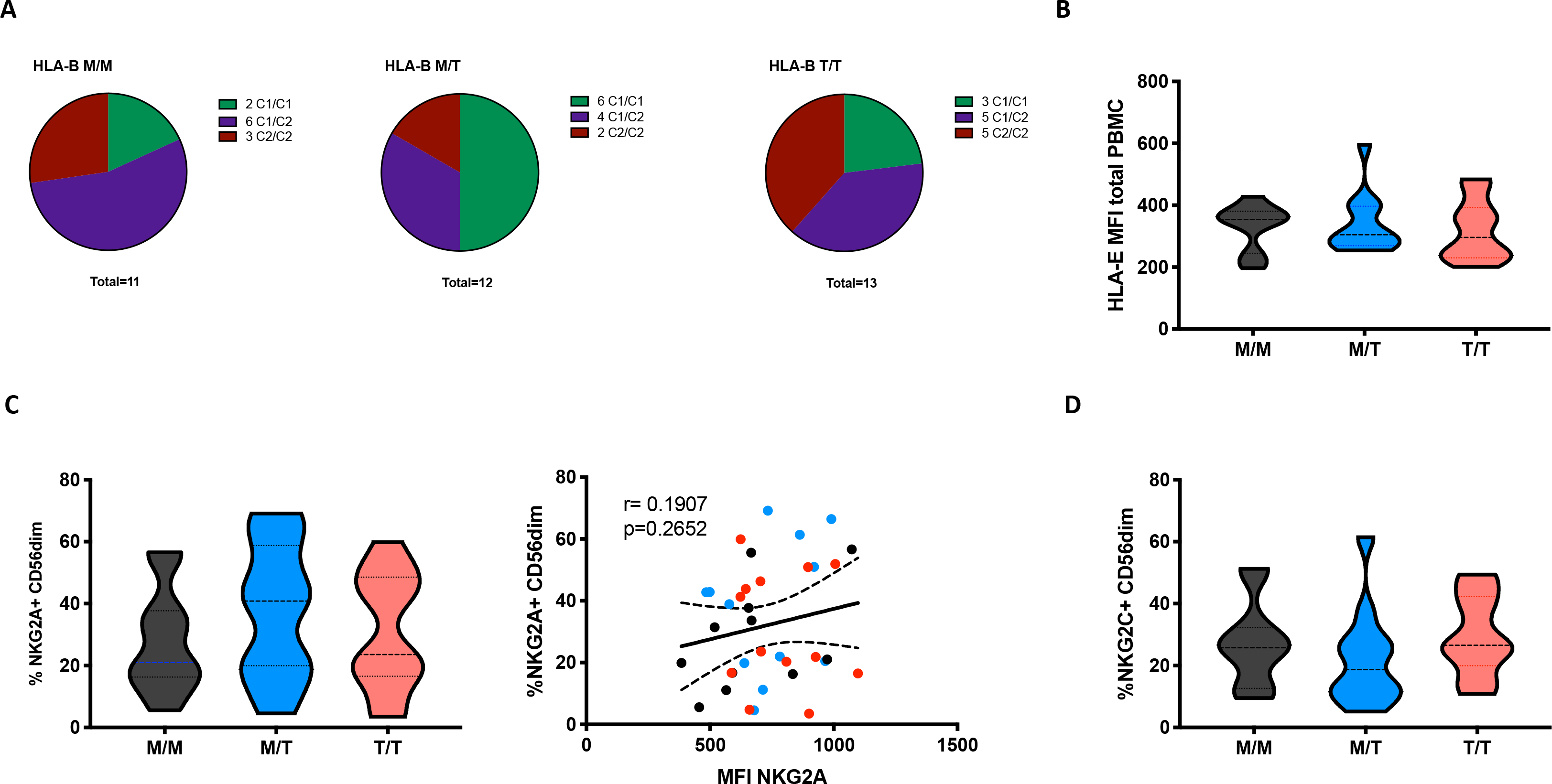
Dimorphism at position −21 HLA-B does not significantly modulate HLA-E and NKG2A expression. (A) HLA haplotypes encoding HLA-C1 and C2 within groups of −21 HLA-B genotype M/M homozygous −21M/T heterozygous and −21 T/T homozygous subjects from the study cohort. (B) Comparison of cell-surface HLA-E expression (MFI of staining with HLA-E-specific antibody 3D12) on total PBMC between groups. Data are displayed as violin plots; the group median and interquartile range are indicated. (C) Propo r tion of CD56^dim^ NK cells (Live CD56+CD3-CD19-CD14-CD4-PBMC) expressing NKG2Abetween groups and compari son of the percentage of NKG2A+ CD56^dim^ with their level of surface expression (MFI) in M/M (black dots), M/T (blue dots) and T/T (red dots) subjects. (D) Violin plots of the frequency of NKG2C expressing CD56_dim_ NK cells in the donor groups. Group median and interquartile range are indicated. All donors were HCMV positive.

To investigate the effects of the HLA-B −21 dimorphism on surface expression of HLA-E we examined the expression of HLA-E on peripheral blood mononuclear cells (PBMCs) in subject groups distinguished on the basis of the amino acid encoded (M/M, M/T and T/T). Despite median levels of total cellular HLA-E expression on PBMCs being higher in M/M individuals, no significant difference was observed in surface HLA-E expression on PBMC between the groups **(Fig. 1B)**. Moreover, no correlation was detected between −21M copy number and the proportion of NK cells expressing either NKG2A **(Fig. 1C)** or its activating counterpart NKG2C **(Fig. 1D)**. These observations contrast findings in HCMV seronegative Eurasian individuals where HLA haplotypes defined by −21M HLA-B were associated with increased surface expression of HLA-E and a decreased frequency of NK cell s expressing NKG2A (6).

Furthermore, no relationship was detected between the surface levels of HLA-E expression and HLA-A imputed expression level (z score) in subject groups distinguished on the basis of the presence of HLA-B −21M in this cohort **(Suppl. Fig. 1A)**. The effect of HLA-A expression on HIV-1 viraemia was also not evident in individuals with HLA-B −21M/M, in keeping with a reported less prominent effect in Africans/African-Americans relative to Caucasians, conceivably as a consequence of the distinct HLA haplotypes and frequencies present in individuals of African descent (Suppl. **Fig.1B)**(Ramsuran et al 2018).

### Surface abundance of HLA-C does not correlate with −21 HLA-B

The amount of cell-surface HLA-C expression has been previously reported to vary with −21 HLA-B type, with M/M HCMV seronegative European donors displaying low levels, as a result of their relatively restricted HLA-C diversity and genotypes dominated by HLA-C*07, which is subject to microRNA-148a (miR-148a) mediated downregulation (18). Whilst in predominantly Eurasian populations haplotypes combining −21M HLA-B and C2 are rare, the African haplotypes present in the M/M group combine specific HLA-B and C2 alleles i.e. B*42:01-C*17:01 and 8*81:01 - C*18:01. Despite some variation in the levels of surface expression of HLA-C (assessed by staining with the HLA-C and HLA-E-reactive antibody DT9) (19), especially in T/T donors, there was no overall difference in the mean levels of expression between the study groups and there was no correlation between cell surface abundance of HLA-C and −21 HLA-B genotype **(Fig. 2A)**. Equally we observed no obvious clustering according to HLA-C1 and C2 types and only a single M/T donor was homozygous for HLA-C*07, an allele that is highly represented in M/M individuals of European origin as previously shown **(Fig. 2B,C**) (6).

**Fig. 2.**
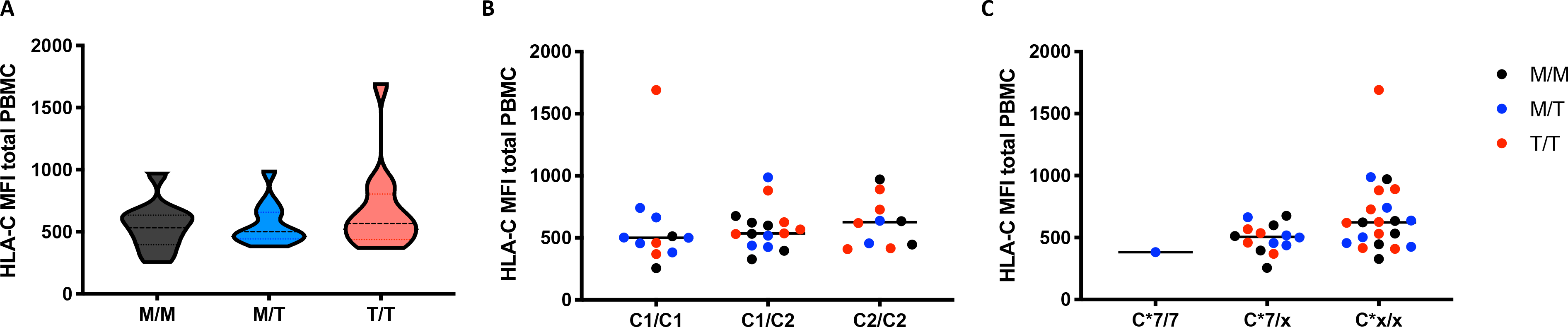
No significant differences in HLA-C expression levels in subjects grouped according to −21 HLA-B dimorphism. Level s of surface expression of HLA-C (MF I of staining with antibody DT-9) on total PBMC in donors grouped: (A) by HLA-B −21 variant (data are shown as violin plots, and group median and quartiles are indicated); (B) according to HLA-C1 and C2 epitopes and (C) by the presence and absence of (x) of HLA-C*07. In panel s (B) and (C), M/M subjects are shown as black dots, M/T as blu e dots and T/T as red dots.

### KIR education and differentiation predominate in HIV/HCMV seropositive subjects of African descent irrespective of −21 HLA-B dimorphism

The effect of viraemic HIV-1 infection on driving alterations in the NK cell subset distribution is well described (20) (15). In keeping with this we confirmed the presence of the aberrant CD56^neg^CD16+NK cell subset in our cohort; however, no significant differences in the frequencies of the CD56^bright^, CD56^dim^ and CD56^neg^ NK cell subsets were observed between groups of M/ M, M/T and T/T individuals (**Suppl Fig. 2 A, B, C)**. Notably chronic HIV/HCMV co-infection also leads to an accentuated differentiation within the CD56^dim^ subset with the emergence of a CD57+NKG2C+KIR+NKG2A- signature and expansion of adaptive NK cell subsets (15) (21) (22). We therefore investigated the phenotypic diversity of CD56^dim^ NK cell subset to delineate the fingerprint of HCMV co-infection in the three study groups in relation to the presence of −21HLA-B dimorphism. In the cohort as a whole, the acquisition of inhibitory KIRs on CD56^dim^ NK cells was paralleled by a loss of NKG2A expression (r=−0.4165, p=0.0143) (23). A tight positive correlation between NKG2C and KIR expression was further noted, in line with expansion of self-specific KIRs in the context of HCMV infection/re-activation (r=0.5960, p=0.0002) (24) (25) (26). The level of expression of KIRs (cocktail of antibodies against KIR2DL1/S5, KIR2DL2/L3/S2, KIR3DL2,and KIR3DL1)on CD56^dim^ NK cells did not differ between the three study groups **(Fig. 3A, B)**. Levels of CD57 expression were also comparable between the three groups, suggesting the presence of NK cells at different differentiation stages **(Fig. 3A, B)**. Examination of additional markers such as the key signalling molecule FcєRl-γ and the transcription factor promyelocytic leukemia zinc (PLZF), the absence of which characterises adaptive NK cell subsets, showed a broader range of expression with a trend for a lower median level of expression in T/T subjects, which did not reach statistical significance **(Fig. 3A, B)**.

**Fig. 3.**
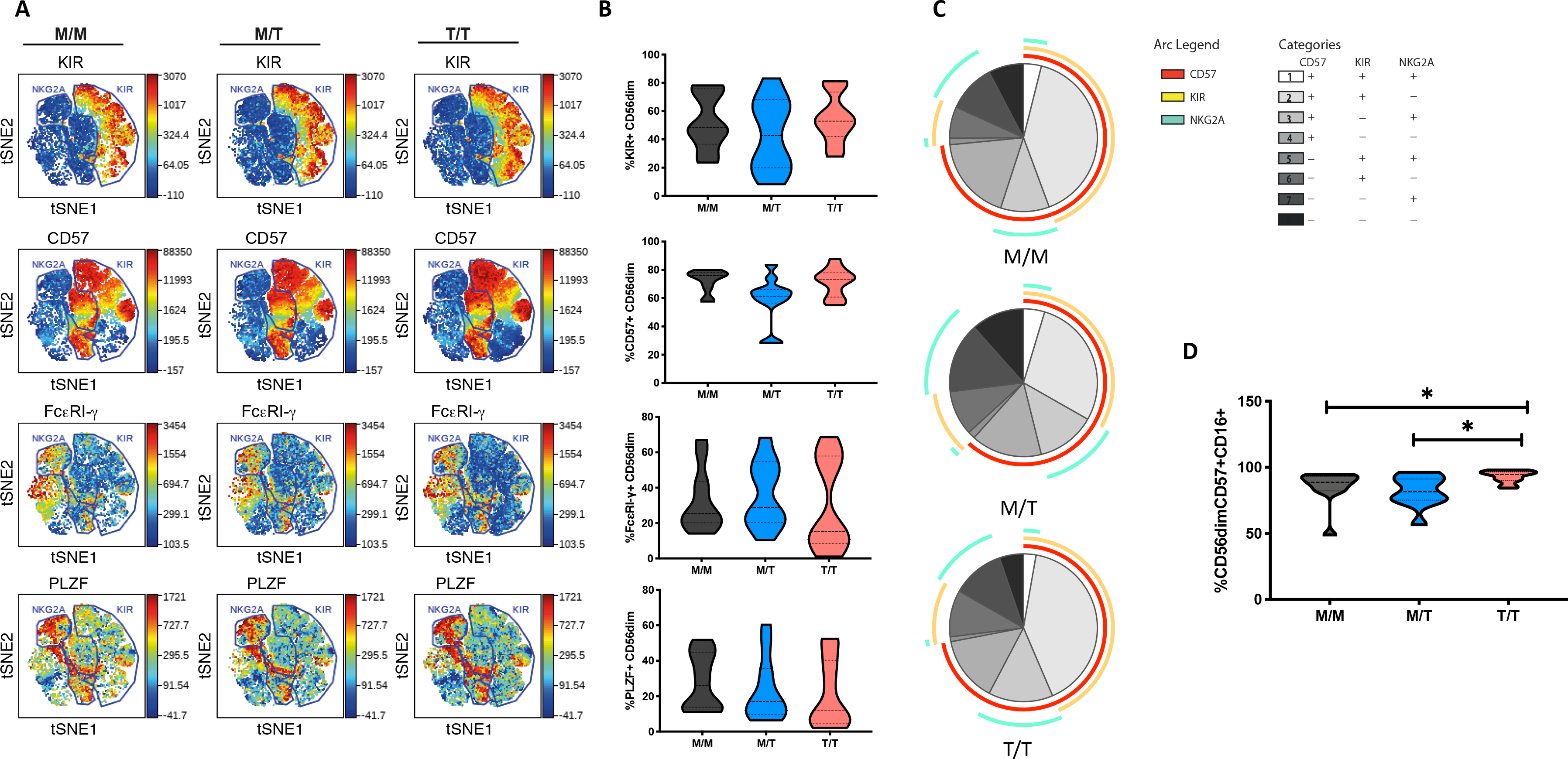
Lack of a dominant effect of −21M HLA-B on the extent of NKG2A driven NK cell differentiation. (A) ViSNE analysis of multiparametric flow data was per formed on CD56^dim^ NK cells from the compiled M/M, M/Tand T/T donors showing expression of KIRs, CD57, FcєRl-γ and PLZF. Each point on the VisNE map represents a single cell and colour depicts intensity of protein expression. (B) Summary violin plots of the proportion of CD56drm NK cells expressing KIRs, CD57, FcєRl-γ and PLZF b etw een donor groups. (C) SPICE analysis pie charts for each group. The pie slices represent the proportion of CD56^dim^ NK cell s expressing different receptorcombin ations, and the pie arcs depict expression of individual receptors, as detailed in the key. (D) Summary violin plots showing the proportion of CD57+CD56^dim^ NK cells expressing CD16 between the donor groups.

Boolean gating analysis was performed next to examine the proportion of NKG2A or KIR-educated CD56^dim^ NK cells that were more highly differentiated (assessed on the basis of expression of CD57) **(Fig 3C)**. The proportion of KIR-NKG2A+ CD56^dim^ NK cells, educated via the inhibitory NKG2A receptor, was 21.95% ±4.432 (mean± *SEM*) in M/M, 29.09% ± 5.315 in M/T and 26.26% ± 5.432 in T/T donors. The extent to which these NKG2A-educated cells were differentiated (according to the expression of CD57) did not vary with the −21M copy number and represented a small fraction (10.65%±2.440, *mean±SEM* in M/M donors, 12.89%± 3.262 in M/T and 14.12%± 3.508 in T/T donors). KIR+NKG2A- NK cells, which can only be educated via KIRs, represented a larger fraction of CD56^dim^ NK cells in all groups(45.83% ±6.159, 36.57% ± 6.847 and 49.48%±5.767, in M/M, M/T and T/T subjects respectively). The proportion of KIR-educated NK cells that were differentiated (CD57+) was higher than that observed in the NKG2A-educated fraction, comprising in M/ M donors 40.43% ± 5.204 in M/M donors and 40.82± 5.339 in T/T donors, and trending to be somewhat lower in M/ T subjects (28.59± 5.629) **(Fig. 3C)**. These results are in keeping with loss of NKG2A expression with increasing NK cell differentiation in HIV infection and contrast findings of a dominant effect of the −21M HLA-B dimorphism on increasing the differentiated subpopulation of the educated KIR-NKG2A+ NK cells in Caucasian HCMV seronegative donors (6).

Whereas all donors exhibited comparable levels of CD57 expression on CD56^dim^ NK cells, the activating receptor CD16 was expressed by a higher proportion of differentiated CD57+ CD56dml NK cells in T/T donors compared to M/M and M/T donors (p=0.02 and p=0.01 respectively) **(Fig. 3D)**. As expected the CD57+ subset of NK cells in all groups was enriched for additional adaptive features such as lower l evels of PLZF and FcєRl-γ and enriched for KIR and NKG2C compared to the CD57 negative fraction of NK cells, as previously described (15). In T/T donors the CD57+ NK cell subset trended to have higher mean levels of expression of NKG2C and lower mean levels of expression of PLZF and FcєRl-γ compared to the CD57+ NK cells in M/M and M/T donors although this did not reach statistical significance(**Suppl. Fig 3**).

### The dominant effect of −21M on NK cell function is lost in HIV/HCMV donors

To further examine the influence of −21 HLA B dimorphism on NK cell education and associated NK cell function we utilised an antibody-coated target cell stimulation assay to measure ADCC. Following stimulation with Raji cells coated with anti-CD20, CD56^dim^ NK cells were assessed for cytokine production by intracellular cytokine staining and degranulation, as measured by surface expression of CD107a. NK cell s from T/T donors demonstrated a trend towards higher production of IFN-γ relative to those from M/M donors and higher IFN-γ production compared to M/T individuals **(Fig. 4A)**. A similar trend towards higher NK cell expression of CD107a was observed in T/T donors in relation to the M/M and M/T groups **(Fig. 4B)**. For both functional responses a range of IFN-γ production and CD107a expression was observed within each group that could not be attributed to their HLA-C haplotype (data not shown). Further analysis of the proportion of IFN-γ producing CD56^dim^ NK cells that were differentiated (according to CD57 expression) and either educated via NKG2A or KIRs, showed a higher proportion of the cytokine producing cells being comprised of CD57+NKG2A-KIR+ NK cells than CD57+NKG2A+KIR- cells in M/M and T/T donors (p= 0.006 and p=0.005 respectively) but this did not reach statistical significance for the M/T group. These differences are reflected in all three pie charts **(Fig. 4C),** where the subset of IFN-γ--producing cells with a CD57+NKG2A-KIR+ phenotype is of similar size in M/M, T/T and slightly smaller in M/T donors, in keeping with lower frequencies of differentiated KIR educated NK cells in this group. These data suggest the dominant effect of KIR mediated education for IFN-γ producing NK cells irrespective of −21 HLA-8 dimorphism. A similar effect was observed for CD107a production in M/M and T/T subjects whereas in M/T donors the effects of the educating KIRs are less distinct (data not shown).

**Fig. 4.**
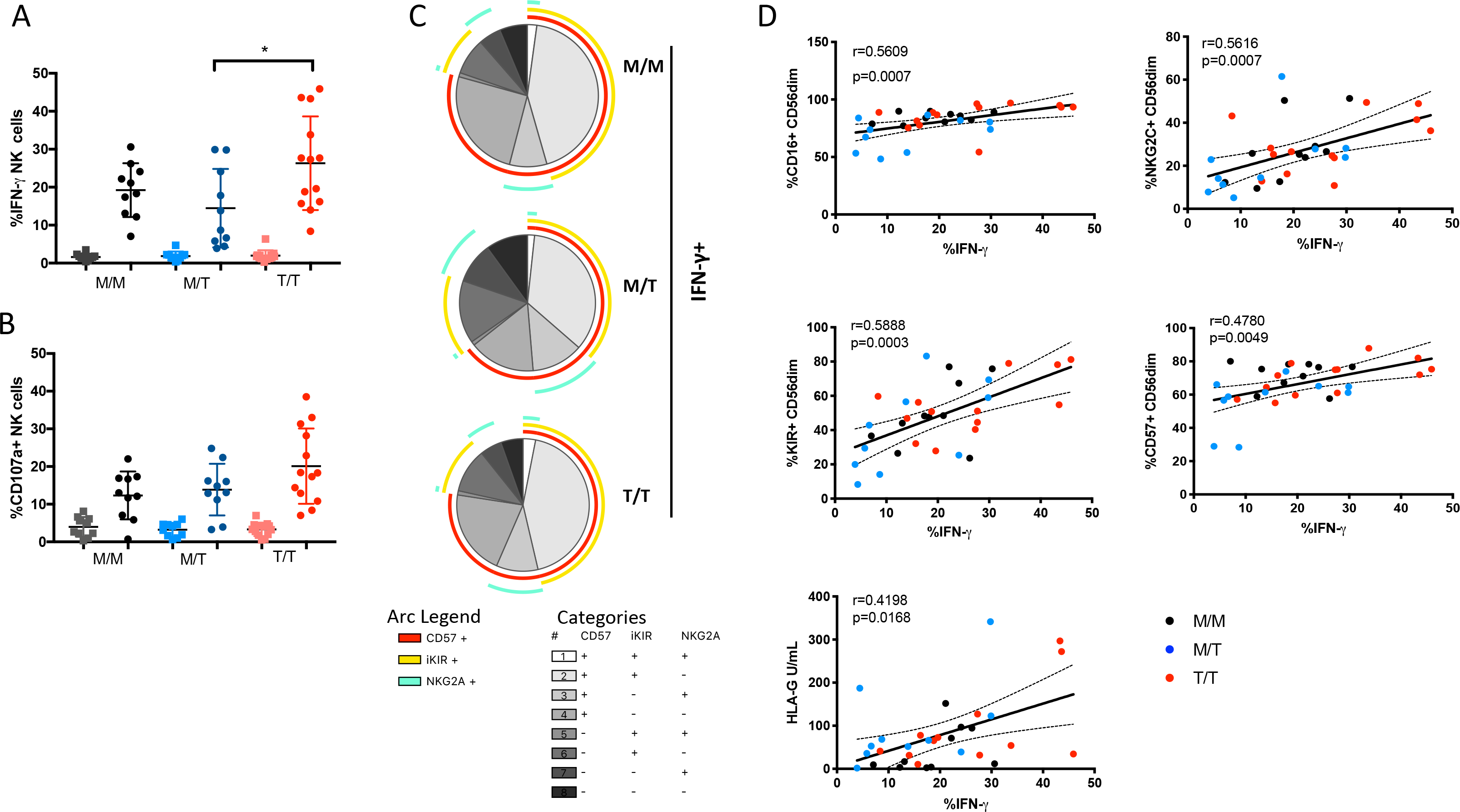
Variable influence of −21 HLA-B dimorphism on ADCC responses. (A) IFN-γ and (B) CD107a expression by CD56drm NK cells foll owing co-culture with RAJI cells coated with anti-CD20 (filled circles) or murine lgG (fill ed squares) in M/M (n=l0), M/T (n=l0) and T/T donors (n=13) wit h available PBMC. (C) SPICE pie charts for each group. The pie slices correspond to the proportion of IFN-γ producing cells that express different receptor combinations, and the pie arcsdepict individual expression of CD57, KIRs and NKG2A, as detailed in the key. (D) Corr elations between IFN-γ production by CD56^dim^ NK cells and expression of CD16, CD57, KlRs, NKG2C or soluble HLA-G levels. The nonparametric Spearman test was used for correlation analysis.

In addition to the effect of education, the variability in the ADCC functional responses of CD56dim **NK** cells, in particular IFN-γ production, could relate to cellular expression of CD16. *Ex vivo* the proportion of NK cells expressnig CD16 correlated with IFN-γ production (r=0.5609 p=0.0007), suggesting that shedding of CD16 reported in progressive HIV-1 infection, could contribute to the reduction of NK cell ADCC function in our study cohort **(Fig. 4D)**(27). Notably adaptive NKG2C+ NK cell subpopulations that arise in response to HCMV infection and expand during HIV-1 infection are imbued with enhanced ADCC capacity, in particular production of IFN-γ following CD16 ligation, reflecting epigenetic modifications and enhanced downstream signalling through CD3z homodimers in the absence of FcєRl-γ (15) (28). We therefore assessed whether the size of adaptive NKG2C+ NK cell populations could account for the variability in IFN-γ production noted between and with in the three study groups. In the cohort as a whole, IFN-γ production correlated strongly (r=0.5616, p=0.0007) with NKG2C expression, suggesting that the presence of adaptive subpopulations, enriched within differentiated CD57+ and KIR+ NK cells (which also correlate with IFN-γ production), could modulate NK cell functional capacity to antibody coated targets **(Fig. 4D)**.

Recently it was demonstrated that HCMV-derived peptides presented by HLA-E, in particular the rare UL40 peptide VMAPRTLFL which is identical to the HLA-G leader peptide, fine tune the ADCC response of NK cells via NKG2C recognition (17). Both membrane bound and soluble levels of HLA-G are reported to be increased in untreated HIV infection and during HCMV infection and have been shown to correlate with blood IFN-γ concentrations and could therefore represent a source of HLA-E peptides in T/T individuals (29) (30) (31). Although we did not detect any significant differences in the soluble plasma HLA-G concentration between the study groups, HLA-G levels showed a weak associaiton with IFN-γ production suggesting that an environment potentially displaying altered HLA-E peptide ligands recognised by adaptive NKG2C expressing NK cells may induce differential cellular responses**(Fig. 4D)**.

## Discussion

HLA-E acts as powerful modulator of the immune response, serving as a ligand for NKG2 receptors that provide a functionally complementary axis to the polymorphic KIR system for control of innate lymphocyte subsets. HLA-E binds signal peptides derived from the leader sequence of HLA-A, B, C and G proteins in order to achieve stable expression at the cell surface (32). −21M, the residue present in all HLA-A and −C and a minority of −B allotypes, facilitates folding and expression of HLA-E by providing a strong anchor residue in contrast to −21T, the residue present in the majority of HLA-B allotypes. Whereas this genetic segregation depending on HLA-B dimorphism leads to a binary form of NK cell education and functional responsiveness in HCMV seronegative donors of European origin by either supplying NKG2A or KIR ligands (6), our findings indicate that this is not a generic principle that applies to ethnically diverse cohorts with HIV/HCMV co-infection. The presence of African specific alleles, together with alterations in the HLA-E peptide repertoire due to the availability of peptides derived from other cellular and viral sources that could arise during HIV/HCMV coinfection, trigger the expansion of adaptive NK cells expressing the activating receptor NKG2C with subsequent functional consequences. The lack of −21M expression could thus become redundant in HCMV seropositive individuals where UL40 or HLA-G derived peptides may stabilise the expression of HLA-E and fine tune NK cell activation and antibody driven adaptive responses.

In Eurasian populations the reported LD between HLA-B −21M and HLA-B Bw6/HLA-C1 limits the supply of KIR ligands and favours NKG2A mediated NK cell education (6). However, the genetic segregation between HLA-C1 and −21M HLA-B was not evident in this study group, where the presence of HLA-C2, a stronger KIR ligand than C1, resulted in the presence of both KIR and HLA-E ligands for NKG2 receptors in M/M donors. In addition, the more common African haplotypes combining −21M and HLA-C2 involve African specific HLA-C allotypes that have leader sequences that poorly promote HLA-E expression, further limiting the supply of HLA-E ligands for interaction with NKG2 receptors on NK cells. These genetic effects could partly explain the lack of association between −21M copy number and surface HLA-E expression in our cohort. Another possible genetic factor that may have influenced the levels of HLA-E expression in our cohort is the dimorphism at position 107 of HLA-E, which distinguishes two most common alleles, HLA-E*0l:01 (position 107 arginine, R) and HLA-E*0l:03 (position 107 glycine, G), the former of which is reported to be expressed at lower levels than the latter (33) although this has not been seen in all studies (9). Nonetheless, although HLA-E genotyping was not performed in our cohort, as the two main HLA-E alleles occur in roughly equal frequencies in different ethnic groups and are maintained in diverse HLA haplotypes by stabilizing selection (34), allele frequencies would not have been expected to differ significantly between our study groups.

In addition to genetic differences between our cohort and those studied previously, the presence of chronic HIV-1 and HCMV co-infection in our study subjects may also have contributed to the lack of significant difference in surface HLA-E expression between study groups. HLA-E surface levels serve as an important sensor of HLA class I expression and are sensitive to perturbations in the biosynthesis of most polymorphic class I allotypes as well as the class Ib molecule HLA-G imparted by viral infections or stress. Of note whilst HIV-1 Nef causes down-regulation of HLA-A, B and Vpu mediates reduction of HLA-C, these viral accessory proteins mediate their effects post-translationally and should not affect the supply of HLA class I signal peptides; and HCMV maintains/stabilizes HLA-E expression (35) (36) (37, 38). However, the presence of specific HCMV UL40 variants and/or HLA-G levels may be altering the supply of HLA-E binding peptides in our cohort.

As well as observing no impact of the HLA-B −21 dimorphism on the level of expression of HLA-E we also did not detect a correlation between −21M copy number and NKG2A expression in our cohort. During NK cell development and education, the acquisition of self-reactive KIRs leads to progressive downregulation and decreased surface expression of NKG2A (23). This process is accelerated during HIV infection/HCMV coinfection and further underlined by the expansion of differentiated CD57+ NKG2C+ NK cell subsets enriched for KIRs for self HLA-C1 and/or C2 allotypes, which explains the lack of correlation between −21M HLA-B and better NKG2A+KIR- educated NK cells in this cohort. Due to limitations in sample availability we were not able to type the KIR genes nor perform staining for individual KIRs in our study subjects. Specific KIR alleles are reported to differ in their strength of signalling, with associated effects on NK cell education/ADCC responses, which could further explain some of the inter-donor variability observed in this study. A recent study has further highlighted the critical role of KIR polymorphism influencing responses to HCMV, where in particular the interaction between KIR2DL1 and HLA-C2 ligands drives large and stable expansions of adaptive NKG2C+ NK cells (26). It would therefore be of interest to determine the effect of KIR polymorphism in modulating the size of the adaptive NK cell pool in larger HIV-1/HCMV coinfected cohorts.

There is increased appreciation that peptides presented via HLA-E during conditions of stress and viral infections influence the activation of NK cells, driving expansion of adaptive NKG2C+ NK cells and subsequent enhancement of ADCC responses. In keeping with this, we observed a range of ADCC responses in our cohort that correlated with NKG2C expression. Furthermore, UL40 in HCMV encodes peptides that mimic MHC class I signal sequences and share a conserved methionine (M) anchor residue at peptide amino-acid residue 2), which correspond to amino acid −21 of the classical HLA class I leader sequence (16) (17). Hence HCMV infection provides peptides that may substitute for host HLA-I-derived HLA-E stabilising nonameric peptides in T/T donors. Interestingly UL40 polymorphisms and the strength of interactions between HLA-E presented peptide and NKG2C controls the activation of adaptive NK cells. Of note, a gradient in NKG2C+ NK cell effector function has been reported depending on the potency of recognition of HCMV peptides (VMAPRTLFL > VMAPRTLIL > VMAPRTLVL) (17). The VMAPRTLFL UL40 derived peptide mimics the signal peptide of HLA-G, the expression of which is upregulated during inflammation, HCMV infection and HIV-1 infection, and specifically enhances antibody-driven adaptive NK cell responses as recently described (17). Regardless of the peptide source, it is tempting to speculate that alterations in the HLA-E ligandome surveyed by the NKG2C receptor contribute differentially to the accumulation, differentiation and effector functions of adaptive NKG2C+ NK cells during infection. Whether HIV-1 peptides could further exploit the HLA-E/NKG2 axis as recently suggested (39) requires further evaluation.

The role of adaptive NK cells in influencing the rate of HIV acquisition and levels of viral control during established infection remains poorly defined, but offers an alternative explanation for previous epidemiological observations, that needs to be formally addressed with a combination of population and functional studies. Assessment of the overall impact of HLA-B dimorphism on the acquisition of or control of HIV-1 infection will need to take into account a number of effects in addition to the contribution of HLA/peptide complex availability and its impact on the NKG2 pathway. These include effects on CD8 T cell responses and interactions between the Bw4/Bw6 epitope and the KIR3DL1/3DS1 pathway, which will necessitate study of much larger cohorts.

In summary, we posit that in addition to differences in the genetic background, chronic HIV-1 infection with frequent reactivations of HCMV affects the pool of peptides presented by HLA-E and surface levels of HLA-E providing a more diverse range of ligands for CD94/NKG2 NK cells. The strength of these interactions and presence of inflammatory stimuli shape the NK cell pool and functional activity, blurring the dichotomous effect of −21 HLA-B on NK cell function seen in Eurasian HCMV seronegative donors. Future larger studies aimed at dissecting the effect of different HLA-E/peptide ligands on adaptive NK cells in relation to −21 HLA-B polymorphism, during disease are required, in order to facilitate realisation of the translational potential of specific NK cell subpopulations and exploit the NKG2C/HLA-E axis to enhance NK cell functionality.

## Author Contributions

EMC performed experiments, contributed to study design, acquisition of data, analysis, and drafting of the manuscript; AO performed experiments and contributed to acquisition of data; PB contributed to study design, data interpretation and critical editing of the manuscript. DP: conception and design of study, data analysis and interpretation, critical revision of the manuscript and study supervision.

## Conflict of Interest

The authors declare that the research was conducted in the absence of any commercial or financial relationships that could be construed as a potential conflict of interest.

## Funding

This work was supported by MRC grants MR/M008614 (DP) and MR/K012037 (PB), an AOP grant awarded to DP and NIH, NIAID, DAIDS UMl grants Al67854 (CHAVI), AI00645 (Duke CHAVI-ID)and AI144371 (BFH, PB) and R56 award AI147778 (PB, DP). PB is a Jenner Institute Investigator.

## Supporting Information

**Supplementary Table 1**. Cohort characteristics and HLA genotypes.

**S1. Suppl. Fig. 1 The effect of HLA-A expression on surface HLA-E expression and HIV viral load (VL) by HLA-B −21 variant**. (A) Surface expression levels of HLA-E (MFI) on total PBMC, according to HLA-A (z score) and HLA-B dimorphism in the study cohort. (B) Correlation of HLA-A expression levels (z score) with HIV VL in HLA-B −21M/M and T/T donors.

**S2. Suppl. Fig. 2 NK cell subset redistribution**. Summary box and violin plots of the frequencies of (A) CD56^bright^, (B) CD56^dim^ and (C) CD56^neg^ NK cell subsets among M/M, M/T and T/T donors. NK cells were gated on live CD-CD14-CD19-CD4- and subsets identified on the basis of CD56 and CD16 expression. Median and interquartile range is shown.

**S3. Suppl. Fig. 3 (A)** Heat map representation of the mean proportion of expression of the markers as shown within the CD56^dim^ CD57+ and CD57-fractions in M/M, M/T and T/T subjects.

