## Supplemental Table 1 for "Subordinate effect of −21M HLA-B dimorphism on NK cell repertoire diversity and function in HIV-1 infected individuals of African origin"

Table 1. Subject characteristics and HLA genenotypes

| Subjects | HLA-B -21 | SEX | Age | Plasma HIV VL (log10) | HLA-B allele 1 | HLA-B allele 1 | HLA-B allele 2 | HLA-B allele 2 | HLA-C allele 1 | HLA-C allele 1 | HLA-C allele 2 | HLA-C allele 2 | Bw4/Bw6 |
|---|---|---|---|---|---|---|---|---|---|---|---|---|---|
| 01 | M/M | F | 26 | 4.287129621 | *08 | *0801g | *14 | *1401 | *02 | *0210 | *16 | *1601 | Bw6/Bw6 |
| 2 | M/M | F | 21 | 4.276714495 | *42 | *4201 | *42 | *4202 | *17 | *1701 | *17 | *1701 | Bw6/Bw6 |
| 3 | M/M | F | 29 | 4.595848805 | *07 | *0702 | *81 | *8101g | *07 | *0702g | *18 | *1801g | Bw6/Bw6 |
| 4 | M/M | F | 22 | 4.068185862 | *14 | *1402 | *42 | *4202 | *08 | *0802 | *17 | *1701 | Bw6/Bw6 |
| 5 | M/M | F | 31 | 2.650307523 | *81 | *8101g | *07 | *0702g | *07 | *0702g | *18 | *1801g | Bw6/Bw6 |
| 6 | M/M | F | 43 | 3.866287339 | *14 | *1401 | *42 | *420101 | *08 | *080201 | *17 | *1701/02/03 | Bw6/Bw6 |
| 7 | M/M | F | 50 | 4.279027806 | *07 | *0702/44/49N/58/59/6 | *42 | *420101 | *07 | *0702/50/66/74/99/117 | *17 | *1701/02/03 | Bw4/Bw6 |
| 8 | M/M | F | 32 | 5.319641047 | *42 | *42:02 | *48 | *48:05 | *17 | *17:01 | *15 | *15:05/09/29/34 | Bw6/Bw6 |
| 9 | M/M | F | 22 | 4.453348923 | *07 | *0702g | *14 | *1402 | *07 | *0702g | *08 | *0802 | Bw6/Bw6 |
| 10 | M/M | F | 23 | 2.602059991 | *81 | *8101/02 | *07 | *0703/16/37/50/*4806 | *18 | *1801/02 | *04 | *0407/*1801/02 | Bw6/Bw6 |
| 11 | M/M | F | 21 | 4.372359583 | *07 | *0703 | *67 | *6701 | *07 | *0702g | *12 | *1203 | Bw6/Bw6 |
| 12 | T/T | F | 22 | 4.708548365 | *15 | *1503g | *35 | *3501/28 | *02 | *0210 | *04 | *0401g | Bw6/Bw6 |
| 13 | T/T | F | 31 | 4.237392915 | *41 | *4101 | *45 | *4501g | *06 | *0602 | *17 | *1701 | Bw6/Bw6 |
| 14 | T/T | F | 19 | 5.238046103 | *15 | *1503g | *45 | *4501g | *02 | *0210 | *06 | *0602 | Bw6/Bw6 |
| 15 | T/T | F | 22 | 4.459392488 | *18 | *1801g | *35 | *3501g | *04 | *04:01/09N/28/30/41 | *07 | *07:04/11 | Bw6/Bw6 |
| 16 | T/T | F | 39 | 3.696356389 | *15 | *1510 | *15 | *1510 | *03 | *0304 | *04 | *0401g | Bw6/Bw6 |
| 17 | T/T | F | 22 | 3.821971818 | *15 | *151001 | *18 | *1801/17N | *07 | *0704/11 | *08 | *0804 | Bw6/Bw6 |
| 18 | T/T | F | 35 | 3.785614525 | *15 | *1503/103 | *45 | *4501/07 | *07 | *0701/06/18/52/57/116 | *16 | *160101 | Bw6/Bw6 |
| 19 | T/T | F | 20 | 4.245191764 | *45 | *4501/07 | *57 | *570301 | *16 | *160101 | *18 | *1801/02 | **Bw4/Bw6** |
| 20 | T/T | F | 35 | 4.479877498 | *44 | *4403 | *45 | *4501g | *14 | *1403 | *16 | *1601 | **Bw4/Bw6** |
| 21 | T/T | F | 29 | 5.459476926 | *18 | *18:01/53/55/59 | *53 | *53:01 | *04 | *04:01/09N/28/30/41 | *07 | *07:04/11 | **Bw4/Bw6** |
| 22 | T/T | F | 23 | 4.460972974 | *18 | *18:01/53/55/59 | *53 | *53:01 | *04 | *04:01/09N/28/30/41 | *07 | *07:04/11 | **Bw4/Bw6** |
| 23 | T/T | F | 28 | 4.678582107 | *15 | *15:03/61/103/173 | *41 | *41:02/11 | *17 | *17:01 | *02 | *02:10 | Bw6/Bw6 |
| 24 | T/T | F | 27 | 2.602059991 | *41 | *4102/04 | *15 | *1503/61/74/*9503 | *02 | *0202/04/07-11/13-15 | *02 | *0203/16/*1701-04 | Bw6/Bw6 |
| 25 | M/T | F | 34 | 4.255586049 | *15 | *1503g | *42 | *4201 | *02 | *0210 | *17 | *1701 | Bw6/Bw6 |
| 26 | M/T | F | 22 | 4.646746713 | *08 | *0801g | *15 | *1510 | *03 | *0304 | *07 | *0701g | Bw6/Bw6 |
| 27 | M/T | F | 31 | 6.019894829 | *14 | *1402 | *18 | *1801g | *07 | *0704/12 | *08 | *0802 | Bw6/Bw6 |
| 28 | M/T | F | 24 | 5.103338401 | *07 | *07:02/44/49N/58/59/ | *18 | *18:01/53/55/59 | *07 | *07:04/11/139 | *07 | *07:02/50/66/74/100/102/ | Bw6/Bw6 |
| 29 | M/T | F | 30 | 4.307496038 | *15 | *1510 | *39 | *3910 | *08 | *0804 | *12 | *1203 | Bw6/Bw6 |
| 30 | M/T | F | 22 | 3.941014244 | *08 | *0801g | *15 | *1510 | *04 | *0401g | *07 | *0701g | Bw6/Bw6 |
| 31 | M/T | F | 21 | 3.739651444 | *15 | *151001 | *08 | *0801/08N/19N | *07 | *0701/06/18/52/57/116 | *08 | *0804 | Bw6/Bw6 |
| 32 | M/T | F | 38 | 3.155943018 | *08 | *0801g | *15 | *1510 | *03 | *0304 | *03 | *0304 | Bw6/Bw6 |
| 33 | M/T | F | 53 | 3.59117595 | *15 | *1503/103 | *39 | *3924 | *02 | *0210 | *07 | *0701/06/18/52/57/116 | Bw6/Bw6 |
| 34 | M/T | F | 27 | 5.170000473 | *15 | *1503g | *42 | *4201 | *02 | *0210 | *17 | *1701 | Bw6/Bw6 |
| 35 | M/T | F | 26 | 4.858687551 | *15 | *151001 | *42 | *4202 | *03 | *030402 | *17 | *1701/02/03 | Bw6/Bw6 |
| 36 | M/T | F | 26 | 4.985821516 | *14 | *14:02/20/22/24 | *15 | *15:10 | *04 | *04:01/09N/28/30/41/57 | *16 | *16:01/32/38/39/41/44 | Bw6/Bw6 |

Suppl. Table 1. Cohort characteristics and HLA genotypes.
