## Supplemental Figure 1 for "Subordinate effect of −21M HLA-B dimorphism on NK cell repertoire diversity and function in HIV-1 infected individuals of African origin"

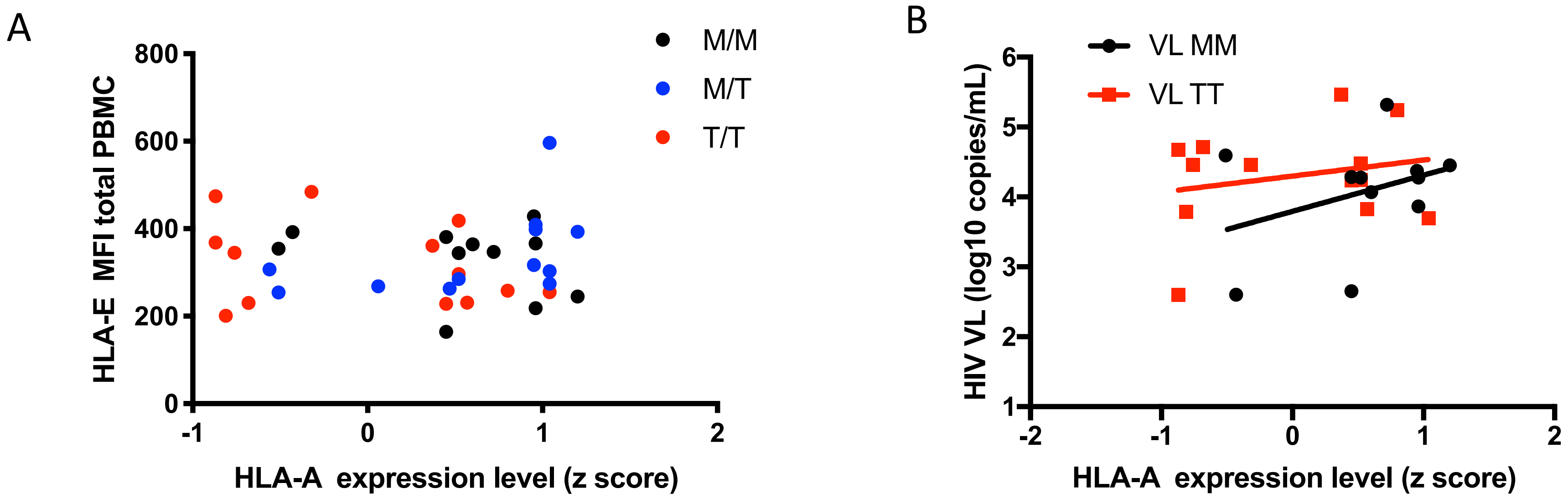


Suppl. Fig. 1 The effect of HLA-A expression on surface HLA-E expression and HIV viral load (VL) by HLA-B -21 variant. (A) Surface expression levels of HLA-E (MFI) on total PBMC, according to HLA-A (z score) and HLA-B dimorphism in the study cohort. (B) Correlation of HLA-A expression levels (z score) with HIV VL in HLA-B -21M/M and T/T donors.
