## Supplemental Figure 2 for "Subordinate effect of −21M HLA-B dimorphism on NK cell repertoire diversity and function in HIV-1 infected individuals of African origin"

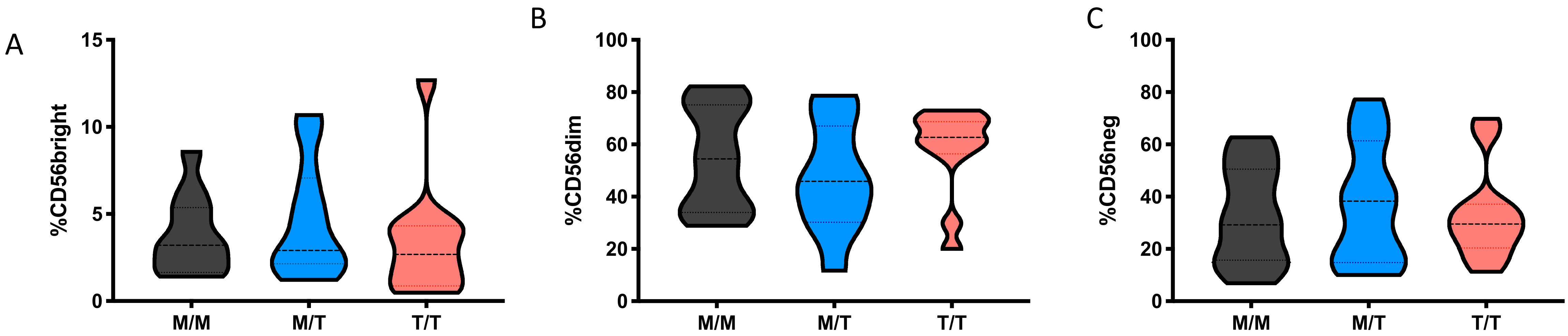


**Suppl. Fig. 2 NK cell subset redistribution.** Summary box and violin plots of the frequencies of (A) CD56$^{bright,}$ (B) CD56$^{dim}$ and (C) CD56$^{neg}$ NK cell subsets among M/M, M/T and T/T donors. NK cells were gated on live CD-CD14-CD19-CD4- and subsets identified on the basis of CD56 and CD16 expression. Median and interquartile range is shown.
