## Supplemental Figure 3 for "Subordinate effect of −21M HLA-B dimorphism on NK cell repertoire diversity and function in HIV-1 infected individuals of African origin"

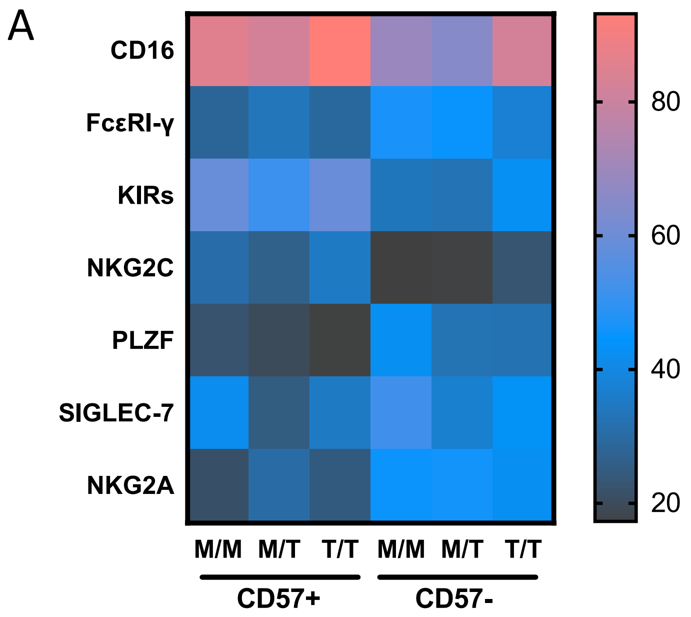


**Suppl. Fig. 3 (A)** Heat map representation of the mean proportion of expression of the markers as shown within the $CD56^{dim}$ CD57+ and CD57- fractions in M/M, M/T and T/T subjects.
